# Neuropeptide Y deficiency in the bone marrow drives hematopoietic stem and progenitor cell aging

**DOI:** 10.64898/2026.02.20.706987

**Authors:** Dinisha Kamble, James P. Ropa, Malgorzata M. Kamocka, Yan Qi, Nick Imperiale, Pratibha Singh

**Affiliations:** Herman B. Wells Center for Pediatric Research, Department of Pediatrics, Indiana University School of Medicine, Indianapolis, IN 46202; Department of Medical & Molecular Genetics, Indiana University School of Medicine, Indianapolis, IN 46202; Indiana Center for Biological Microscopy, Department of Medicine, Indiana University School of Medicine, Indianapolis, IN 46202

## Abstract

Aging-related blood disorders are linked to defects in the regenerative and multilineage differentiation ability of hematopoietic stem and progenitor cells (HSPCs). While remodeling of the bone marrow (BM) microenvironment where HSPCs reside is known to contribute to these age-associated defects, the underlying factors and mechanisms remain poorly defined. Here, we discovered that the age-related decline of the neurotransmitter neuropeptide Y (NPY) in the BM is a critical driver of HSPC dysfunction. Using mouse models, we demonstrated that NPY levels decrease in the BM with age, and that genetic NPY overexpression or exogenous NPY administration in old mice substantially reverses aging-associated phenotypic and functional defects in HSPCs. Transcriptome analysis revealed that NPY supplementation in old mice restores aging-disrupted molecular pathways in their HSCs, including oxidative stress responses, myeloid differentiation, stemness, mitochondrial activity, and RhoA signaling. However, NPY genetic loss in young mice led to a decline in HSCs regenerative capacity and increased oxidative stress. Importantly, NPY levels also decline in elderly humans, and ex vivo treatment of human BM-derived HSPCs with NPY enhances their in vivo repopulating capacity. These results suggest that NPY supplementation or preservation of NPY-producing nerve fibers could be a therapeutic strategy to rejuvenate aged HSC function.

## Introduction

The global elderly population is rapidly increasing, and aging is a major risk factor for hematologic disorders. Aging reduces the regenerative ability of hematopoietic stem and progenitor cells (HSPCs) and biases their differentiation towards the myeloid lineage (1). These defects in HSPCs contribute to aging-related blood disorders, including increased susceptibility to infection, risk of anemia, and predisposition to myeloid neoplasms (2). Age-associated dysfunction of HSPCs has been linked to alterations in the bone marrow (BM) microenvironment/also known as niche, where HSPCs primarily reside and are essential for maintaining their function (3–5). However, the factors and mechanisms involved are not completely understood. The nerve fibers innervate the BM, and neurotransmitters released by these nerve fibers critically regulate HSPC function (3, 6–9). Recent studies indicate that changes in adrenergic signals in the BM with aging contribute to HSPC defects (3, 7), but the role of other neurotransmitters remains unexplored. We previously reported that neuropeptide Y (NPY), a neurotransmitter produced in the BM primarily by the sympathetic nerve fibers, critically regulates HSPC trafficking by controlling the BM vascular gateway function (9). NPY is also shown to promote hematopoietic regeneration after genotoxic stress (10). Accumulating evidence suggests a connection between NPY and longevity, as NPY-deficient mice are reported to have reduced lifespan benefits of dietary restriction (11), whereas NPY overexpression in rats extends lifespan (12). Additionally, lower circulating NPY levels were found in elderly human populations than in younger individuals (13). However, it is currently unknown whether aging affects local NPY levels in the BM and if these changes contribute to age-associated defects in HSPCs. Using genetic and pharmacological approaches in mouse models, we identify NPY deficiency in the BM as a causal driver of HSPC aging. We found that NPY levels decline in mouse BM during aging, and genetic NPY overexpression or exogenous NPY administration in old mice substantially alleviates aging-related phenotypic and functional abnormalities in HSPCs. Conversely, NPY loss in young mice induces multiple hallmarks of HSPC aging. RNA-sequencing reveals that NPY supplementation reverses age-associated molecular pathways linked to myeloid differentiation, redox regulation, mitochondrial activity, and RhoA signaling. Ex vivo culture studies demonstrate that NPY improves clonal proliferative capacity of aged mice HSPCs by suppressing the RhoA-ROCK1 pathway. Together, these findings establish declining BM NPY as a key niche-derived regulator of HSC aging and identify NPY as a potential strategy to rejuvenate aged hematopoiesis.

## Results

### NPY levels decline with age, and genetic overexpression of NPY in mice mitigates age-associated defects in HSPCs

Aging-associated degradation of adrenergic nerve fibers and alteration in adrenergic signals in the BM have been linked to a decline in HSPC function (3, 7, 14). However, the effects of aging on other neurotransmitter levels in the BM and their potential contributions to HSPC aging are not well understood. We previously reported that NPY plays a vital role in G-CSF-induced HSPC egress from BM to circulation (9). During a search for the neurotransmitters that might change during aging, we measured NPY levels in the BM of young (3-4 months) and old (18-20 months) wild-type (WT) mice. Compared to young WT mice, old WT mice showed reduced NPY levels in the BM extracellular fluid (Figure 1A). To evaluate if aging impacts NPY levels in humans, we measured NPY levels in the plasma samples of young and elderly humans. Older individuals (55-73 years) exhibited lower NPY levels compared to younger individuals (30-45 years) (Figure 1B), which was consistent with a previous report (13). Aging leads to several phenotypic and functional defects in HSPCs, including increased numbers, decreased repopulation ability, and a shift towards myeloid differentiation (3, 5, 15, 16). Reduced levels of NPY in elderly humans and aged mice prompted us to investigate whether NPY deficiency contributes to the decline in HSPC function associated with aging. We suggested if aging-related NPY deficiency is involved in HSPC defects, then NPY overexpression should alleviate HSPC defects in old mice. To explore this possibility, we used WT young (3-4 months) and 18-20 months old WT and NPY overexpressing (NPY OE; heterozygous) mice. Analysis of HSPC subpopulations in the BM of these mice revealed that, aligned with previous reports (3, 15, 16), WT old mice showed a substantial increase in HPC enriched Lineage-Sca-1+c-Kit+(LSK) frequencies and HSCs enriched Lineage-Sca-1+c-Kit+CD150+CD48-(SLAM LSK) (Supplementary Figures 1A &1B) and numbers (Figure 1C-1E) compared to WT young mice. However, aging-related increases in SLAM LSK and LSK observed in WT old mice were attenuated in NPY OE old mice (Figure 1C-1E; Supplementary Figures 1A &1B). SLAM LSK counts also increased in the spleen of WT old mice compared to WT young mice, but not in NPY OE old mice (Figure 1F). HPC1 (Lineage^-^Sca-1^+^c-Kit^+^CD150^+^CD48^+^) and HPC2 (Lineage^-^Sca-1^+^c-Kit^+^CD150^-^CD48^+^) frequencies were similar in these three groups of mice (Supplementary Figures 1C-1E). Multipotent progenitor cells (MPP; Lineage^-^Sca-1^+^c-Kit^+^CD150^-^CD48^-^) frequencies were higher in WT old mice than in WT young mice, and this aging-related increase was substantially attenuated in NPY OE old mice (Supplementary Figures 1C & 1F). Mature myeloid cells frequencies increased and lymphoid cells frequencies decreased in WT old mice compared to WT young mice (Figure 1G), that was consistent to previous reports (17, 18). However, the aging-related increase in myeloid cell frequencies was attenuated in NPY OE old mice, but lymphoid cells were not restored (Figure 1G). Total cell counts in the BM and spleen were comparable among young, WT old, and NPY OE old mice (Figure 1H & 1I). In the PB of the three groups of mice, similar WBCs, lymphocytes, and neutrophils were observed (Supplementary Figures 1G-1I). Monocytes showed a trend toward increased counts, while platelet counts were significantly higher in WT old mice than in their young counterparts. However, NPY OE old mice did not exhibit this increase in monocytes or platelets (Supplementary Figures 1J-1K). Importantly, the age-related decline in BM NPY levels observed in WT old mice was abolished in NPY-OE old mice, which maintained young-level NPY expression (Figure 1J). Tyrosine hydroxylase (TH) staining, used as a surrogate marker of nerve integrity (19), showed reduced TH signals in the BM of WT old mice compared with young mice, whereas this decline was attenuated in NPY-OE old mice (Supplementary Fig. 1L-1M). This data suggests that age-related reductions in NPY likely stem from nerve damage, and that NPY overexpression mitigates this damage. We next evaluated whether NPY overexpression can improve aging-associated defects in HSPCs function. BM cells from WT old mice had lower HSPC clonal proliferation than those from WT young mice, but NPY OE old mice BM did not (Figure 1K). We assessed the repopulation capacity of HSCs from WT young, WT old, and NPY OE old mice by a competitive transplantation. Purified 500 SLAM LSK cells from these mice along with 0.5 million competitor whole BM cells were transplanted into lethally irradiated recipients (Figure 1L). In line with previous studies (3, 14), HSCs from WT old mice exhibited a marked decline in hematopoietic repopulation compared with WT young HSCs (Figure 1M). Importantly, this decline was attenuated in HSCs from NPY OE old mice (Figure 1M). Multilineage differentiation analysis showed that HSCs from WT old mice were skewed toward myeloid differentiation at the expense of lymphoid differentiation (Figure 1N). Whereas NPY OE old mice HSCs exhibited a differentiation pattern resembling that of young mice (Figure 1N). Six months post-transplantation, BM analysis of recipients showed a higher frequency of WT old HSCs than WT young HSCs (Figure 1O), resembling the aged steady-state profile (Figure 1E). However, NPY OE HSC frequencies remained similar to those of young HSCs (Figure 1O). To further validate the repopulation capacity of donor HSCs, we performed secondary transplantation. WT old mice HSCs showed substantially diminished engraftment compared to young mice HSCs, whereas this decline was alleviated in old NPY-OE HSCs (Figure 1P & 1Q). Of note, WT young and NPY OE young mice exhibited comparable frequencies of LSK, SLAM LSK, HPC1, HPC2, and MPP subpopulations in the BM, along with similar BM cellularity and HSPC repopulation capacity (Supplementary Figure 2A-2G).

**Figure 1.**
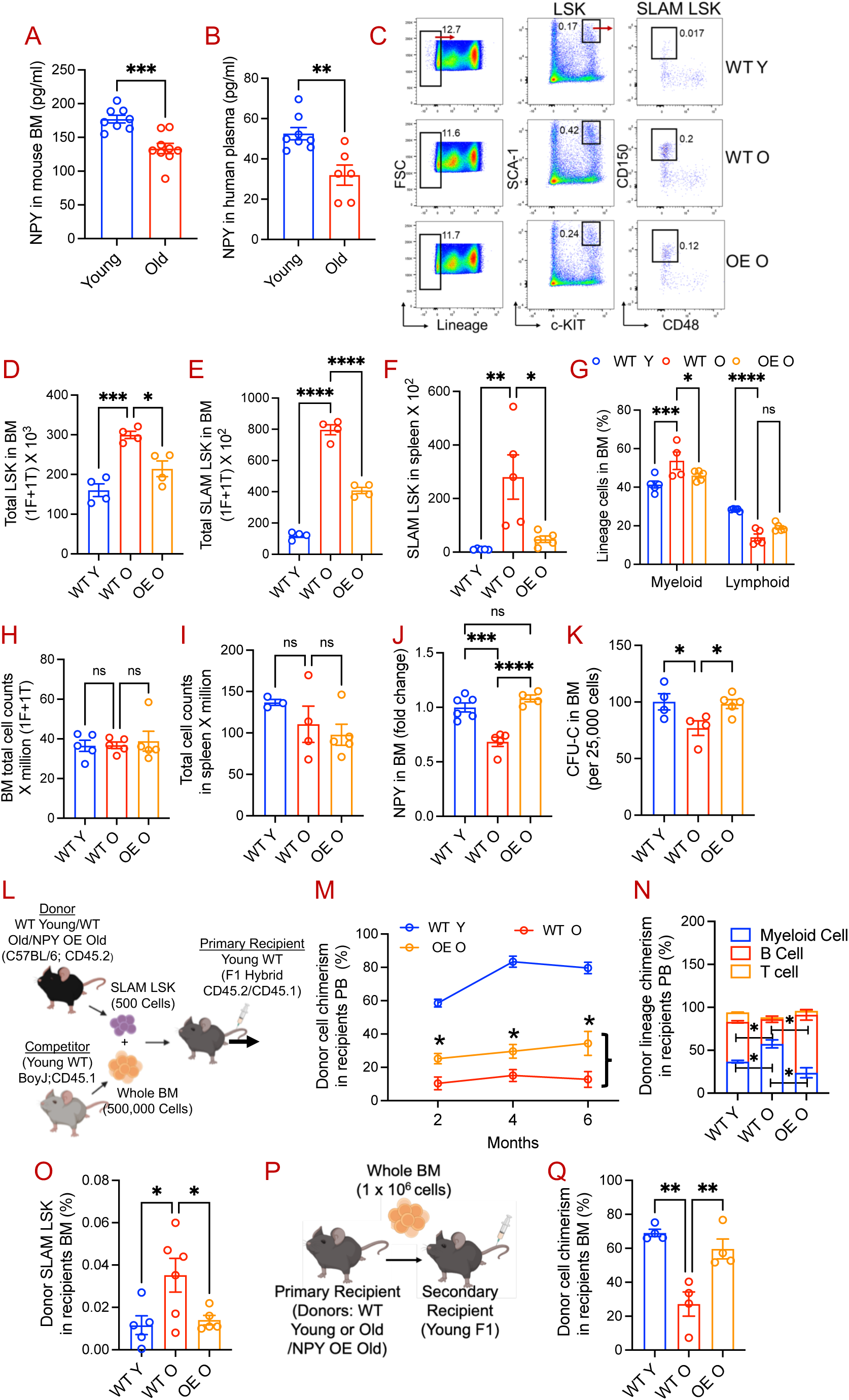
NPY overexpression in mice mitigates age-associated decline in HSPC repopulation capacity. **(A)** NPY levels in the BM of young (3-4 months) and old (18-20 months) mice, and **(B)** NPY levels in plasma from young (30-35 years) and old (55-73 years) human donors, measured by ELISA. **(C-E)** Flow cytometry gating strategies for phenotypic HPCs (Lin^-^c-Kit^+^Sca-1^+^; LSK) and HSCs (Lin^-^c-Kit^+^Sca-1^+^CD48^-^CD150^+^; SLAM LSK), and their quantification in the BM of WT young, WT old, and NPY OE old mice. **(F)** HSC counts in the spleen. **(G)** Myeloid and lymphoid cell populations in the BM assessed by flow cytometry. **(H-I)** Total cellularity of the BM and spleen measured using a Heska analyzer. **(J)** NPY levels in the BM of WT young, WT old, and NPY OE old mice. **(K)** Ex vivo clonogenic proliferation of HPCs assessed by CFU-C assays. **(L-O)** Evaluation of HSC repopulation and lineage differentiation in primary competitive transplantation. **(L)** Schematic of the competitive transplantation experiment. BM SLAM LSK cells (500 cells) from WT young, WT old, or NPY OE old mice were mixed with 0.5 million whole BM competitor cells and transplanted into lethally irradiated recipient mice. **(M)** Donor-derived chimerism in PB at 2, 4, and 6 months post-transplantation. **(N)** Donor-derived myeloid and lymphoid lineage contributions in PB at 6 months post-transplantation. **(O)** Donor-derived HSCs in the BM of recipient mice at 6 months post-transplantation. **(P-Q)** Evaluation of HSC repopulation capacity in secondary transplantation **(P)** Schematic of secondary transplantation. One million BM cells from primary recipients (6 months post-transplantation) were transplanted into lethally irradiated secondary recipients. **(Q)** Donor-derived chimerism in the BM of secondary recipients assessed at 3 months post-transplantation. Data are presented as mean ± S.E.M.; N=4-7 mice; *p ≤ 0.05, **p ≤ 0.01, ***p ≤ 0.001.

We next evaluated the effects of NPY overexpression on previously reported HSC-intrinsic aging hallmarks (15, 20, 21). As reported (22), CD61+ myeloid-biased HSCs were higher in the BM of WT old mice than in WT young controls. Notably, this age-associated increase was diminished in NPY OE old mice (Figure 2A). These observations align with our findings that NPY overexpression mitigates aging-induced myeloid-biased differentiation (Figures 1F and 1M). Consistent with previous reports (23, 24), HSCs from WT old mice exhibited increased cell cycling (Figure 2B), elevated ROS levels (Figure 2C), reduced MMP (Figure 2D), and increased DNA damage (Figure 2E; Supplementary Figure 2H) compared with HSCs from young mice. Notably, these aging-associated changes were substantially alleviated in HSCs from NPY OE old mice (Figure 2B-2E). Given that age-associated changes in the BM microenvironment contribute to HSPC dysfunction, we evaluated BM niche cell populations (Figure 2F). WT old mice exhibited significantly reduced endothelial cells (ECs; CD45^-^Ter119^-^CD31^hi^VE-Cadherin^+^) and mesenchymal stromal cells (MSCs; CD45^-^Ter119^-^CD31^-^CD51^+^PDGFRa^+^) compared with young WT controls (Figure 2G & 2H). These age-related declines in EC and MSC numbers were markedly attenuated in NPY OE old mice (Figure 2G & 2H). HSC-supporting factors like SDF-1, VEGF A and VEGF D were also reduced in the BMEF of WT old mice but remained intact in NPY OE mice (Supplementary Figure 2I-2K). Moreover, inflammatory factors, including IP-10, IL-1b, MIG, MIB-1b, and G-CSF, were elevated in the BMEF of WT old mice but were preserved in NPY OE old mice (Supplementary Figure 2L-2P). Overall, these data suggest that genetic NPY overexpression alleviates aging-related defects in the HSC and BM niche components.

**Figure 2.**
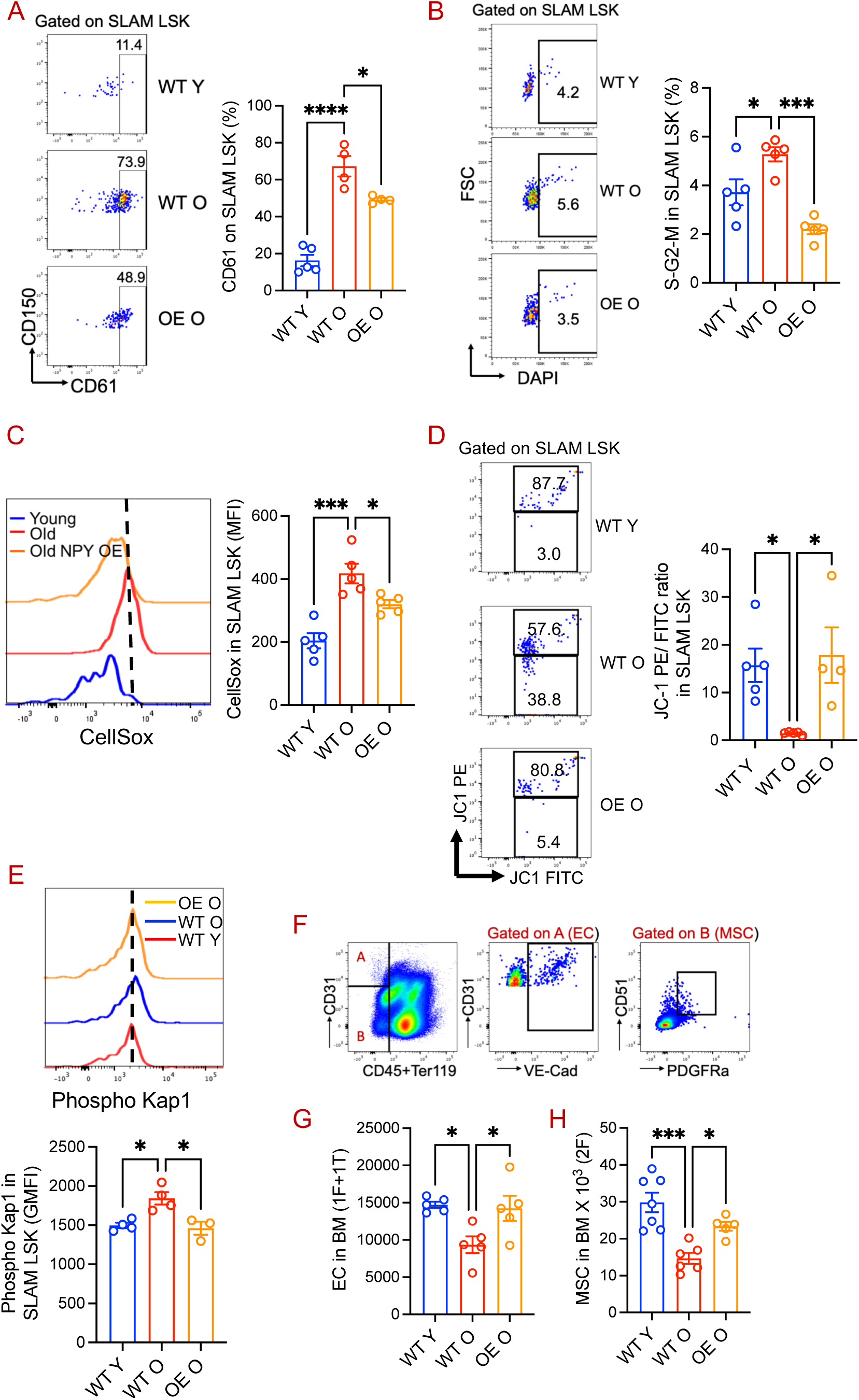
NPY overexpression in mice attenuates aging-associated alterations in HSCs and the BM niche. BM HSCs from WT young, WT old, and NPY OE old mice were evaluated to measure: **(A)** CD61 expression, **(B)** cell-cycle by DAPI staining, **(C)** ROS production by CellROX staining, **(D)** MMP assessed by JC-1, and **(E)** DNA damage by phospho Kap1. **(F-H)** The BM niche was examined by analyzing ECs and MSCs: **(F)** representative flow cytometry gating strategy for ECs and MSCs; **(G)** quantification of ECs; and **(H)** quantification of MSCs in the BM. Data are presented as mean ± S.E.M. (n = 4-7 mice per group). *p ≤ 0.05; ***p ≤ 0.001.

### Pharmacological NPY supplementation in old mice reverses HSC aging

To evaluate the therapeutic relevance of NPY in aged HSC rejuvenation, we utilized a pharmacological approach and administered exogenous NPY in WT old mice for 15 constitutive days (100 ng/mouse). Similar to the NPY OE old mice (Figure 1C-1E), NPY supplementation in WT old mice substantially reversed aging-related increase in LSK and SLAM LSK frequencies (Supplementary Fig. 3A-3B) and their counts in the BM (Figure 3A & 3B). Total BM cellularity was similar in these mice (Figure 1C). The frequencies of HPC1 and HPC2 subpopulations were similar in young, old, and NPY-treated old mice, while the age-related increase in MMPs was reversed by NPY treatment (Supplementary Fig. 3C-3E). CMP and GMP proportions were elevated in untreated old mice but normalized with NPY treatment (Supplementary Fig. 3F & 3G), while MEP frequencies were unchanged across groups; CLPs declined with aging and were not restored by NPY (Supplementary Fig. 3H & 3I). NPY administration in old mice improved the ex vivo clonal expansion capacity of their HPCs (Figure 3D). In competitive transplantation assays, purified HSCs (500 SLAM LSK) from old mice showed a reduced repopulation ability compared to those from young mice (Figure 3E & 3F). However, this age-related decline in HSC repopulation capacity was partially reversed in HSCs from NPY-treated old mice (Figure 3E & 3F). Of note, the expression levels of CXCR4 and CD49d, key adhesion markers involved in HSC homing, were comparable in these mice (Supplementary Fig. 3J & 3K). Multilineage differentiation analysis of repopulated donor cells from young, old, and NPY-treated old mice showed myeloid-biased lineage differentiation from old mice HSCs (Figure 3G). However, this aging-associated myeloid-skewed lineage differentiation was substantially reverted in HSCs from NPY-treated old mice (Figure 3G). Evidence of rejuvenation in NPY-treated old mouse HSCs was also observed in secondary transplantation recipients (Figure 3H-3I). HSCs from NPY-treated old mice exhibited enhanced engraftment and increased lymphoid differentiation relative to those from untreated old mice (Figure 3J). NPY treatment in old mice substantially reversed key aging-related hallmarks in HSCs, including reduced CD61 expression, decreased cell cycling, lower cellular and mitochondrial ROS levels, and a trend toward restored MMP (Figure 3K-3O). Additionally, NPY reverted the aging-related reduction in BM MSC numbers and their clonal expansion (Figure 3P & 3Q) but not ECs (Supplementary Fig. 3L). NPY treatment in aged mice restored age-related declines in HSC-supporting niche factors (SDF-1 and VEGF) in the BMEF, reduced pro-inflammatory factors (MIG, TNF-α, and IP-10), and restored anti-inflammatory cytokine IL-4 (Supplementary Fig. 4A-4G). These results collectively suggested that NPY supplementation in old mice can substantially reverse the aging-related decline in HSC function and BM niche remodeling. To assess whether NPY can improve human HSPC function, we ex vivo cultured BM-derived CD34+ cells from healthy donors across a wide age range (20-47 years old) with or without NPY and transplanted them into sublethally irradiated NSG mice. NPY-treated cells showed improved repopulation compared with untreated controls, regardless of donor age (Figure 3R).

**Figure 3.**
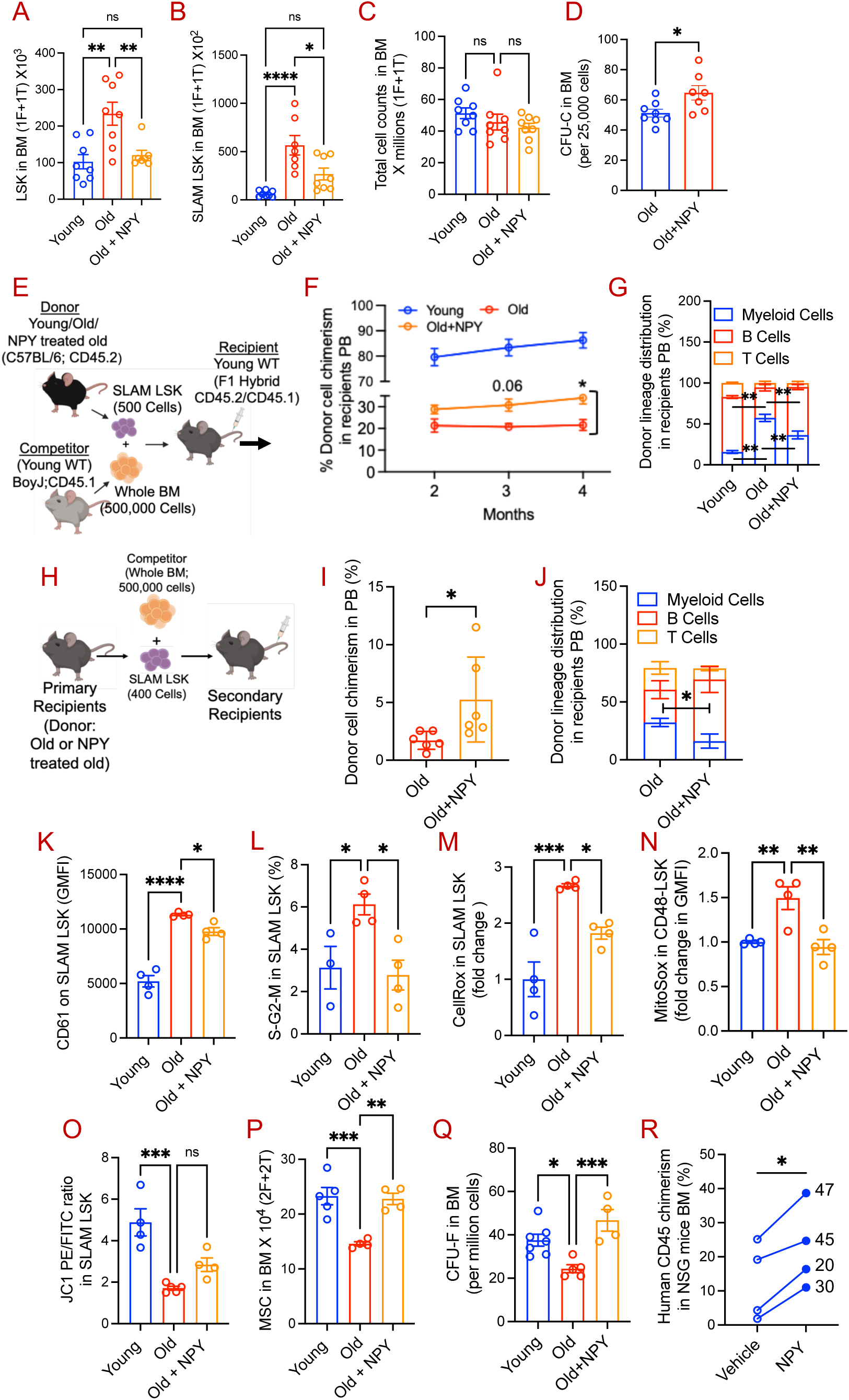
Pharmacological NPY supplementation in WT old mice substantially reverses HSC aging. NPY was administered to WT old mice (100 ng/day, subcutaneously) for 15 consecutive days, and 16-20 hours after the last injection, HSPC and niche cells were evaluated**. (A-B)** Phenotypic HPC and HSC quantitation by flow cytometry in the BM of young, old, and NPY-supplemented old mice. **(C)** BM cellularity measured using Heska analyzer. **(D)** BM HPC clonogenic expansion evaluation by CFU-C assay. **(E-G)** Measurement of hematopoietic repopulation and multilineage differentiation evaluation of HSCs from young, old, and NPY-treated old mice by competitive transplantation. **(E)** Schematic of competitive transplantation. FACS-purified 500 SLAM LSK from these mice were mixed with 0.5 million whole BM competitor cells and transplanted into lethally irradiated recipient mice. **(F)** Donor cell overall chimerism in the PB of recipients at different time points post-transplantation, and **(G)** myeloid and lymphoid distribution in the PB of recipient mic at four months post-transplantation. **(H-J)** Evaluation of donor HSC repopulation by secondary transplantation. (H) Schematic of secondary transplantation. SLAM LSK (400 cells) were purified from the BM of primary transplantation recipients and mixed with 0.5 million whole BM competitor cells. These cells were transplanted into lethally irradiated mice, and (I) donor cell chimerism (J) lineage differentiation in the PB of recipients was evaluated at three months post-secondary transplantation. **(K-O)** Young, old, and NPY-treated old mice BM HSC CD61 expression (K), Cell cycle by DAPI staining (L), cellular ROS by CellRox (M), mitochondrial ROS by MitoSox (N), and MMP by JC-1 staining (O) using flow cytometry. (P) Quantitation of MSC (CD45^-^Ter119^-^CD31^-^PDGFRa^+^CD51^+^) numbers **(Q)**, and evaluation of BM MSC clonal expansion ability in the BM of young, old, and NPY-treated old mice. Data are mean ± S.E.M.; N=4-7 mice; *p ≤ 0.05, **p ≤ 0.01, ***p ≤ 0.001. **(R)** Effect of ex vivo NPY treatment on human BM CD34+ repopulation. Human BM CD34+ cells (human age is shown on the right side in the graph) were cultured for 24 hours with or without NPY (100 ng/mL) in StemSpan medium, then transplanted into sublethally irradiated NSG mice. At three months post-transplantation, human cell chimerism in the BM of recipient mice was evaluated by flow cytometry. Data are mean ± S.E.M.; N=4 human BM samples; *p ≤ 0.05.

### NPY treatment in old mice restores the molecular pathways associated with HSC aging

To identify the molecular pathways involved in NPY-dependent attenuation of HSC aging, we performed RNA sequencing of purified HSCs from young, old, and NPY-treated old mice and compared the transcriptome expression (Figure 4A). Among the DEGs, 1,797 genes were upregulated and 1,549 downregulated in HSCs from old versus young mice (Supplementary Fig. 4H). In NPY-treated old mice, 165 genes were upregulated and 173 downregulated compared with untreated old mice (Supplementary Fig. 4H). Analysis of the unique and shared gene expression using a Venn diagram showed that the expression of 93 genes was restored in NPY-treated old mice HSCs, overlapping with those in HSCs from young mice (Figure 4B). Gene Set Enrichment Analysis (GSEA) revealed that several hallmark pathways related to HSC aging, such as myeloid differentiation, MAPK signaling, immune response, RHO GTPase, and IL-12 signaling, were downregulated in HSCs from NPY-treated old mice compared to those in old mice (Figure 4C).

**Figure 4:**
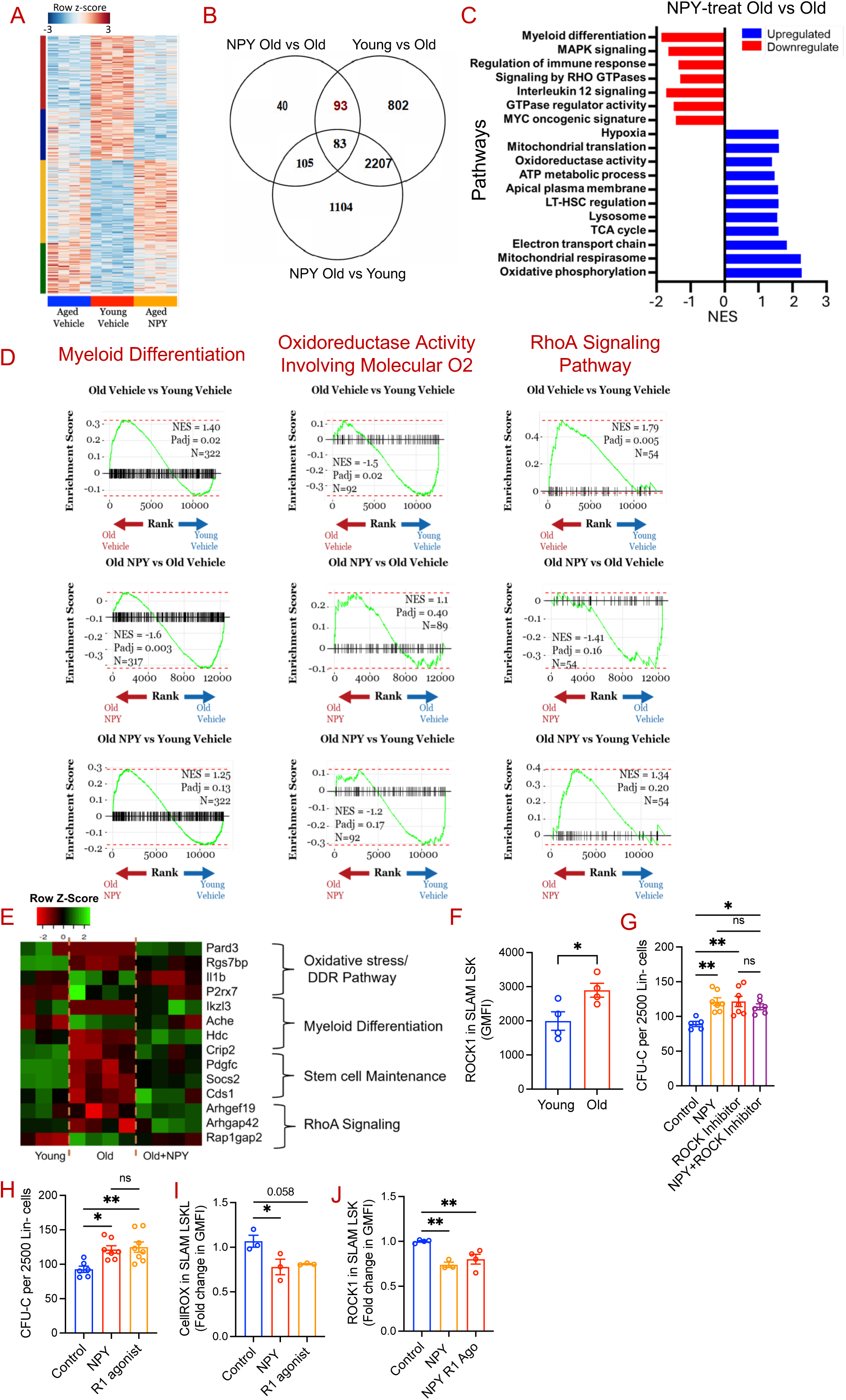
Exogenous NPY administration in WT old mice reverts molecular pathways contributing to HSC aging. **(A)** RNA-seq analysis of HSCs from young, old, and NPY-treated old mice. The Heatmap illustrates the clustering of differentially expressed genes. **(B)** Venn diagram showing unique and shared gene expression in HSCs from young, old and NPY treated old mice analyzed by Venny 2.1. (padj < 0.05). **(C)** GSEA bar graph showing enrichment of molecular pathways in HSCs from NPY treated old vs old mice, and **(D)** Enrichment plots from fgsea for molecular pathways associated with myeloid differentiation, resistance to oxidative stress, RhoA signaling. Data are the Normalized Enrichment Score (NES), adjusted p-value (padj), and number of genes in the data set overlapping with the ranked list (N). **(E)** Heat map of gene expression levels in HSCs from young, old, and NPY-treated mice (n=4 mice/group; Log2FC>1.5; FDR<0.05). **(F)** Evaluation of ROCK1 expression in the BM HSCs of young and old mice (Data are mean ± S.E.M.; N=4 mice; *p ≤ 0.05). **(G)** Effect of NPY treatment and ROCK inhibition on old mice HSPC clonal expansion. Lineage negative BM cells from old mice were cultured in StemSpan medium and treated will NPY (100 ng/mL) or Y-27632 alone (10 µM) or in combination for three days. 2500 ex vivo cultured cells from each culture condition were used to evaluate HSPC proliferation in CFU-C assay. **(H-J)** Effect of NPY R1 receptor agonist treatment on aged mice HSPCs clonal expansion, ROS production, and ROCK1 expression. Lineage-negative BM cells from old mice were cultured with NPY (100 ng/mL) or [Leu31, Pro34]-NPY (1 nM), an NPY Y1 receptor agonist for three days as described above, followed by quantitation of **(H)** HSPC clonal expansion by CFU-C assay, **(I)** ROS production, and **(J)** ROCK1 expression. (Data presented as mean ± S.E.M.; *p ≤ 0.05, **p ≤ 0.01).

Whereas gene sets linked to hypoxia, oxidoreductase activity, long-term HSC regulation, and oxidative phosphorylation were upregulated in NPY-treated mice HSCs (Figure 4C). Myeloid differentiation pathways were markedly increased in old mice HSC but were strongly attenuated in NPY-treated old mice (Figure 4D; left), aligning with our finding that NPY treatment reverts myeloid-skewed lineage differentiation (Figure 3G). Additionally, pathways related to oxidative stress resistance were downregulated in old HSC but partially restored in NPY-treated old HSC, making them comparable to young HSC (Figure 4D; Middle), consistent lower ROS production in NPY treated old mice HSC (Figure 3M). Furthermore, RhoA signaling pathways increased with aging, and NPY treatment partially restored them to levels seen in young HSC (Figure 4D; Right). RhoA signaling pathways are crucial for HSC polarity and engraftment (25, 26). In HSCs from NPY-treated old mice, downregulation of RhoA signaling corresponds with improved hematopoietic engraftment (Figure 3F & 3I). Genes related to oxidative stress, DNA damage response (*Pard3*, *Rgs7bp*, *Il1b*, and *P2rx7*), myeloid differentiation (*Ikzf3*, *AChE*, *Hdc*, and *Crip2*), stemness (*Cds1*, *Socs2*, and *Pdgfc*), and RhoA pathways (*Arhgef19*, *Arhgap42*, and *Rap1gap2)* show impaired expression in old HSCs but were significantly restored in NPY-treated old mice HSCs (Figure 4E). These findings suggest that NPY rejuvenates old HSCs by reprogramming several molecular pathways. Because NPY partly reverses aging-induced upregulation of RhoA pathways, we next investigated the role of the NPY-RhoA axis in alleviating aged HSPC defects. RhoA, a small GTPase, activates Rho-associated coiled-coil-containing kinase (ROCK), a downstream effector that regulates biological activities. We measured ROCK1 and ROCK2 expression in the BM SLAM LSK of young and old mice. HSCs from old mice showed higher ROCK1 levels compared to those from young mice (Figure 4F), while ROCK2 levels remained unchanged (not shown). Because NPY administration in aged mice improves HSPC clonal expansion (Figure 3D), we next examined whether this effect is mediated through modulation of the RhoA/ROCK pathway. Lineage-negative, HSPC-enriched BM cells from old mice were cultured ex vivo with NPY, the ROCK inhibitor Y-27632, or a combination of both. We reasoned that if NPY enhances aged HSPC clonal expansion by inhibiting the RhoA/ROCK pathway, then co-treatment with a ROCK inhibitor would not confer an additional benefit. Indeed, NPY and Y-27632 each increased HSPC clonal expansion to a similar extent, and their combined treatment did not produce further improvement (Figure 4G). NPY exerts its biological effects through at least three G-protein–coupled receptors-R1, R2, and R5 (9)-all of which are expressed on LSK cells from both young and aged mice (Supplementary Figure 4I). To assess whether NPY enhances clonal expansion of aged HSPCs via R1 signaling, lineage-negative BM cells from aged mice were cultured ex vivo with NPY or the selective R1 agonist, [Leu31, Pro34] NPY. Similar to NPY, the R1 agonist improved HSPC clonal expansion (Figure 4H). In addition, treatment with the R1 agonist, like NPY, reduced ROS production and ROCK1 expression in aged HSCs (Figure 4I and 4J). These data suggest that NPY promotes rejuvenation of aged HSPCs through R1-mediated suppression of the RhoA/ROCK pathway.

### Genetic deletion of NPY in young mice triggers premature HSC aging

Because NPY overexpression or supplementation alleviates aging-associated HSC dysfunction, we investigated whether complete NPY loss in young mice induces premature HSC aging. HSPC phenotypes and functions in young NPY KO mice were compared with age-matched WT controls, and old WT mice served as a physiological aging reference. Similar to aged mice, young NPY KO mice displayed increased frequencies and absolute numbers of SLAM LSK cells in both the BM and the spleen compared to young WT mice (Supplementary Fig. 5A; Figure 5A & 5B). However, these increases were less pronounced than those observed in old WT mice. The LSK frequencies and counts were comparable between young WT and KO mice (Supplementary Fig. 5B Figure 5C). Despite this, BM cells from young NPY KO mice formed fewer CFU-C colonies, mirroring age-related functional decline (Figure 5D). Other progenitor subpopulations, including HPC1, showed a downward trend; HPC2 was reduced; and MPPs were elevated in young KO mice compared with WT controls (Supplementary Fig. 5C-5E). Unlike aged mice, which show increased CMP and GMP populations and reduced CLPs (Supplementary Fig. 3F, 3G & 3I), these committed progenitor populations were unchanged between young WT and KO mice (Supplementary Fig. 5F-5I). Analysis of mature hematopoietic populations in the BM revealed a selective reduction in B cell frequencies in young NPY KO mice, similar to that observed with aging, whereas T cell and myeloid cell frequencies remained unaffected (Figure 5E-5G). Total BM and spleen cellularity did not differ between these mice (Supplementary Fig. 5J & 5K). In PB, total WBC, lymphocyte, and neutrophil counts were comparable; however, young NPY KO mice exhibited increased monocyte counts and decreased platelet numbers (Supplementary Fig. 5L-5P). To assess HSC functional competence, 500 SLAM LSK cells from young or old WT or young NPY KO mice were transplanted into lethally irradiated young WT recipients (Figure 5H). HSCs from young NPY KO mice showed significantly reduced long-term repopulation capacity, resembling aged HSCs, although the defect was milder (Figure 5I). Unlike aged HSCs, which exhibit myeloid-biased differentiation, lineage differentiation from young NPY KO HSCs remained balanced and similar to young WT controls (Figure 5J). Expression of homing-associated molecules CXCR4 and CD49d was unchanged, indicating that defective engraftment was not due to impaired homing (Supplementary Fig. 5Q-5R). Notably, the repopulation defect persisted upon secondary transplantation, demonstrating a durable intrinsic HSC impairment (Figures 5K and 5L). To evaluate the contribution of the recipient microenvironment in HSPC aging, we transplanted BM cells isolated from young WT donor mice into middle-aged (12-month-old) WT and NPY KO recipient mice. Three months post-transplantation, donor-derived hematopoietic reconstitution was significantly reduced in the BM of NPY KO recipients compared with WT controls (Supplementary Fig. 5S), suggesting impaired niche support in the absence of NPY. Lineage analysis of donor-derived cells revealed a skewing toward myelopoiesis in NPY KO recipients, characterized by increased frequencies of myeloid cells and a concomitant reduction in B lymphoid cells relative to WT recipients (Supplementary Fig. 5T). Consistent with observations in middle-aged recipients, young NPY KO recipients also exhibited reduced repopulation of young WT donor cells compared with age-matched WT recipients. However, unlike in the middle-aged cohort, the lineage distribution of donor-derived cells was comparable between young WT and KO recipients (data not shown), suggesting that an NPY-deficient aged microenvironment drives myeloid-biased differentiation. Moreover, HSCs isolated from young NPY KO mice demonstrated increased proliferative activity, as evidenced by elevated cell cycle, together with increased cellular and mitochondrial ROS levels (Figure 5M-5O).

**Figure 5:**
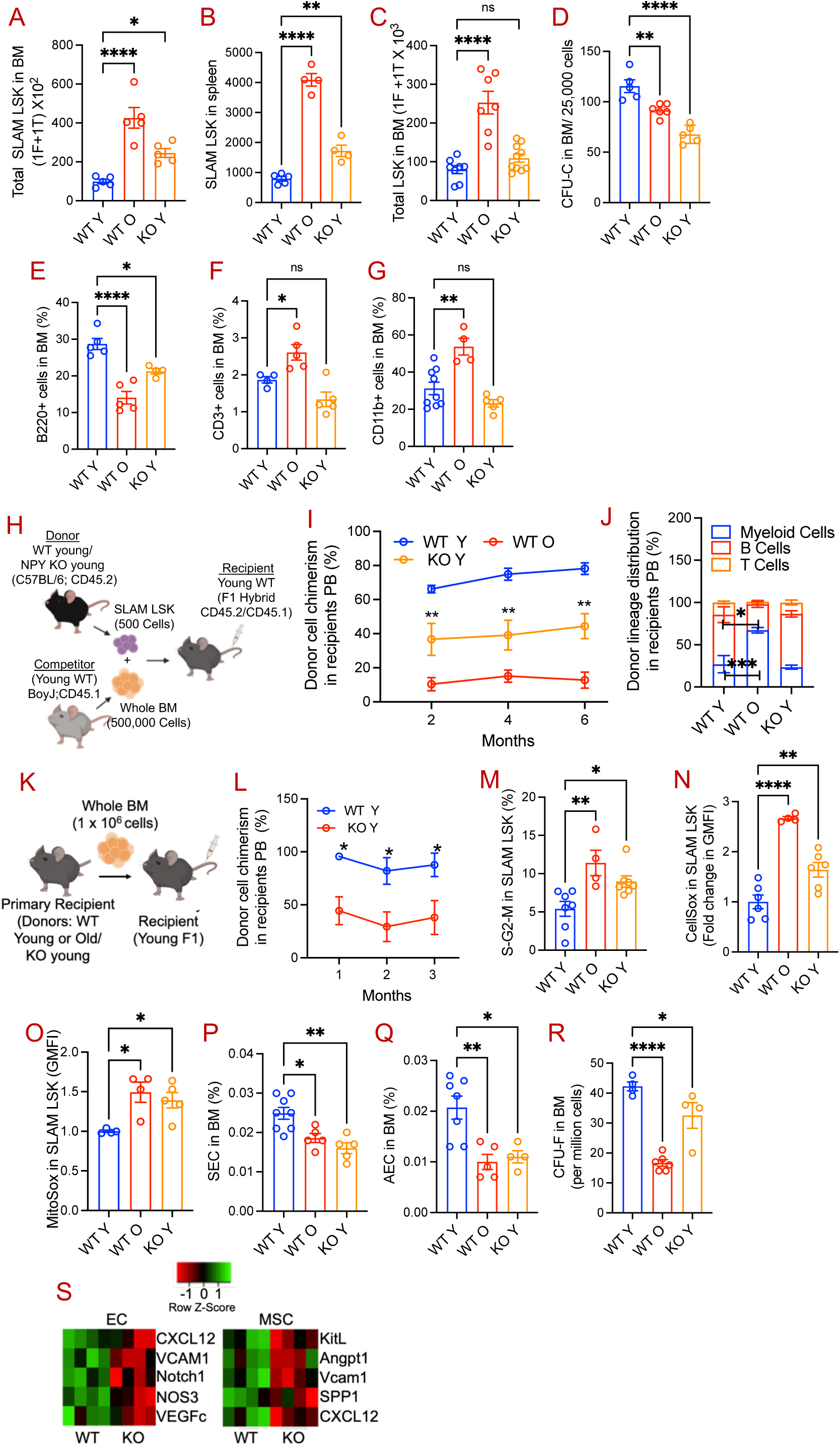
NPY deficiency in young NPY KO mice triggers premature HSC aging. (A-B) Phenotypic HSC counts in the BM and spleen of young and old WT and young NPY KO mice. (C) Phenotypic HPC counts in the BM. (D) Evaluation of HPC clonal proliferation ability by CFU-C assay. (E-G) Mature hematopoietic cells including, (E) B cells, (F) T cells and (G) Myeloid cells in the BM of young and old WT and young NPY KO mice. (H-J) Evaluation of HSC repopulation ability of young and old WT and young NPY KO mice. (H) Schematic of competitive transplantation. FACS-purified 500 SLAM LSK from these mice were mixed with 0.5 million whole BM competitor cells and transplanted into lethally irradiated recipient mice. (I) Donor cell overall chimerism in the PB of recipients at different time points post-transplantation, and (J) myeloid and lymphoid distribution in the PB of recipient mice at six months post-transplantation. (K-L) Evaluation of donor HSC repopulation by secondary transplantation. (K) Schematic of secondary transplantation. (L) Donor cell chimerism in the PB of secondary transplanted recipients. (M-O) Evaluation of HSC properties of young and old WT and young NPY KO mice including, (M) cell cycle, (N) cellular ROS, (O) mitochondrial ROS. (P-Q) Sinusoidal ECs (CD45^-^Ter119^-^CD31^hi^VE-Cad^+^Sca-1^int^) and arteriole ECs (CD45^-^Ter119⁻CD31^hi^VE-Cad⁺Sca-1^hi^) cells in the BM. (R) MSC clonal expansion quantitation by CFU-F assay. (S) HSC-supporting factors transcript expression in the BM EC (CD45^-^Ter119^-^CD31^+^VE-Cadherin^+^) and MSC (CD45-Ter119^-^CD31^-^CD51^+^PDGFRa^+^) of young WT and KO mice. Data are mean ± S.E.M.; N=4-6 mice; *p ≤ 0.05, **p ≤ 0.01, ***p ≤ 0.001.

Analysis of the BM niche revealed that young NPY KO mice had reduced frequencies of sinusoidal ECs (CD45^-^Ter119^-^CD31^hi^VE-Cad^+^Sca-1^int^) and arteriolar ECs (CD45^-^Ter119⁻CD31^hi^VE-Cad⁺Sca-1^hi^), changes similar to those observed during aging (Figures 5P and 5Q). Although total MSC numbers were unchanged (Supplementary Fig. 5U), MSCs from young KO mice showed reduced clonal proliferative capacity, albeit to a lesser extent than aged MSCs (Figures 5R). Furthermore, BM MSCs and ECs from NPY KO mice expressed lower levels of key HSC-maintenance factors (Figure 5S), paralleling transcriptional changes seen in old mice (3, 4). Collectively, these data demonstrate that genetic loss of NPY in young mice is sufficient to induce several abnormalities in HSC and BM niche cells, which are typically observed during physiological aging.

## Discussion

Neuronal signals regulate hematopoiesis through receptors expressed on HSCs and BM niche cells (6, 8, 10, 27, 28). Age-associated damage in BM adrenergic nerve fibers has been linked to HSC dysfunction (3). Here, we demonstrate that NPY levels decline with age, contributing to impaired HSC function. Importantly, both genetic overexpression and exogenous administration of NPY substantially mitigate aging-associated HSC defects, including reduced hematopoietic repopulation capacity, myeloid-biased differentiation, increased cell cycling, and elevated ROS production. Conversely, complete loss of NPY in young mice induces several premature aging phenotypes in HSCs.

Aging-related HSC dysfunction has been linked to inflammation in the BM niche (3, 4, 16, 29–32). In older individuals, increased production of inflammatory cytokines (33) by monocytes and macrophages (18, 34, 35) leads to chronic inflammation, which likely contributes to myeloid-biased lineage differentiation. We found that NPY levels decline with age and that restoring NPY in aged mice reduces inflammatory cytokines and myeloid biased differentiation, suggesting that age-associated loss of neuronal signaling contributes to these defects. Consistent with this finding, NPY has previously been shown to suppress inflammatory cytokine production during bacterial infections (36). Aging is associated with alterations in BM niche components and reduced levels of HSC-supporting factors (3, 4, 16). Notably, NPY alleviates these age-related BM niche abnormalities, suggesting a protective role for NPY in preserving BM niche function during aging.

Aging also affects HSCs through intrinsic changes, including metabolic shifts, oxidative stress, and DNA damage. Young HSCs primarily rely on glycolysis for energy, while aged HSCs switch to oxidative phosphorylation, which increases ROS production (37, 38). This shift, along with impaired antioxidant defenses, leads to oxidative stress, contributing to DNA damage. DNA repair mechanisms are also impaired during aging (39), leading to the accumulation of damaged DNA that increases inflammation due to the senescence-associated secretory phenotype (SASP) (40), as well as an increase in myeloid-biased HSCs (41, 42). Our findings show that genetic overexpression and administration of NPY reduce ROS production and DNA damage in old HSCs while also decreasing the myeloid-biased HSCs. GSEA analysis indicates that HSCs from aged mice treated with NPY exhibit coordinated upregulation of hypoxia-responsive pathways and mitochondrial bioenergetic programs, including oxidative phosphorylation, the TCA cycle, and ATP metabolism. These findings suggest that NPY improves aged HSC function, likely by restoring a metabolically efficient, niche-adapted HSC state with enhanced mitochondrial function. Our findings suggest that NPY improves aged HSC function, likely by both acting directly on HSC and indirectly via niche cells, but further study is needed to determine which is functionally indispensable.

In summary, age-related reductions in NPY drive BM niche remodeling and HSC dysfunction, and restoring NPY signals represents a promising strategy to reverse niche aging and rejuvenate HSC function.

## Methods

Sex as a biological variable. Our study included both males and females, and similar findings were reported in both sexes.

## Mice

10-12 weeks old wild-type (WT) C56BL/6J mice were purchased from Jackson Labs, and Boy J and F1 hybrid mice were from the IUSCC In Vivo Therapeutics Core facility. 14-16 month old WT C57BL/6J mice were received from NIA and aged up to 20 months. NPY knockout (129S-Npytm1Rpa/J) and NPY overexpressing (B6;129S4-Npytm2Rpa/J) mice were purchased from the Jackson Laboratory, backcrossed on C57BL/6J and aged up to twenty months in our facilities. Both females and males were used throughout study. All animal studies were approved by the Indiana University School of Medicine Institutional Animal Care and Use Committee.

### Reagents

NPY and anti-mouse/human NPY ELISA kit from Millipore Sigma (Burlington, MA); Mouse lineage depletion kits from Miltenyi Biotech (Auburn, CA). Antibodies (anti-mouse lineage cocktail, CD117, Sca-1, CD150, CD48, CD11b, CD3, B220, CD51, PDGFRa, CD31, VE-cadherin) from BioLegend (San Diego, CA). CellROX, MitoSOX, LIVE/DEAD staining kit, Tetramethylrhodamine Methyl Ester (TMRM) kit and collagenase Type I from Thermo Fisher Scientific (Waltham, MA. Anti-phospho KAP1 from Abcam (Waltham, MA); anti-NPY 1R from Santa Cruz (Dallas, TX); anti-ROCK1 from R&D Systems (Minneapolis, MN); Leu31 Pro34-NPY from Alpha Diagnostics (San Antonio, TX). MethoCult™ GF, MesenCult Expansion media, and Y-27632 dihydrochloride from STEMCELLS Technologies (Vancouver, BC, Canada). BIBO-3304 trifluoroacetate from Cayman Chemicals (Ann Arbor, Michigan).

### Hematopoietic and non-hematopoietic cell isolation

Mice were sacrificed using CO2 inhalation followed by cervical dislocation. BM cells were harvested by crushing femurs and tibias in cold α-modified Eagle medium (α-MEM) (Lonza Biologics, Walkersville, MD) with 2% fetal bovine serum (FBS) (Corning, Glendale, Arizona). To isolate MSC and ECs, crushed femurs and tibias were incubated with 0.25% collagenase type-1 in 20% FBS for 15 min at 37 °C, and cells were filtered through 40 µm filters. Blood samples were obtained via cardiac bleeding.

### Hematopoietic stem transplantation

HSC-enriched SLAM LSK (Lineage^-^Sca1^+^c-Kit^+^CD48^-^CD150^+^) from the BM of young or old WT, NPY KO, and NPY OE mice (all are C57BL/6 background; CD45.2) were sorted using FACS. These cells were mixed with 0.5 million whole BM cells from Boy J mice (CD45.1) and transplanted into lethally irradiated (11 GY, split dose) young C57BL/6 X Boy J F1 hybrid (F1; 3-4 months old) mice. Donor cell chimerism in peripheral blood (PB) was evaluated up to six months post-transplantation. For secondary transplantation, 1 x 10^6^ whole BM cells or purified SLAM LSK of primary transplantation recipients were transplanted into lethally irradiated young F1 hybrid mice. To examine the effect of the recipient’s niche on the hematopoietic repopulation of donor cells, young WT mice-derived BM cells (one million) were transplanted into lethally irradiated middle-aged (12 months old) WT or NPY KO mice. Three months post-transplantation, donor cells’ hematopoietic repopulation and lineage differentiation were evaluated in the BM of recipient mice.

### HPC Colony Assay

To evaluate HPC ex vivo clonal proliferation, 25000 whole BM cells or 2500 lineage depleted BM cells were cultured in MethoCult™ GF M3434 and incubated in a humidified atmosphere of 5% CO2 and 5% O2 for 7 days. After culture, the colonies were counted manually using an Inverted light Microscope (ECHO REBEL) at 40x magnification.

### Ex Vivo culture of HSPC-enriched BM cells

Lineage-negative BM cells from 18-month-old WT mice were cultured in StemSpan medium supplemented with SCF, TPO, and Flt3 ligand (20 ng/mL each) and treated for 3 days with NPY (100 ng/mL), a ROCK inhibitor Y-27632 (10 µM), either alone or in combination, or with an NPY Y1 receptor agonist [Leu31, Pro34]-NPY (1 nM). Cells were then used for CFU-C quantitation (2,500 cells plated) and for measuring ROS production and ROCK1 expression.

### ELISA

Bone marrow extracellular fluid (BMEF) was collected by crushing femurs and tibias using a mortar and pestle in 1 mL of ice-cold PBS, followed by centrifugation at 4000 rpm for 5 minutes. These BM soups were used to measure NPY levels using an ELISA kit.

### RNA sequencing

Total RNA was extracted from 4000-6000 sorted HSCs using the RNAeasy Plus Micro kit (Qiagen) and assessed for integrity with an Agilent 2100 Bioanalyzer (Agilent Technologies). Library preparation and sequencing were performed at The Center for Medical Genomics, Indiana University, using the NovaSeq 6000 (Illumina). Quality control was conducted with FastQC, and reads were mapped to the reference genome using STAR. The RNA-seq data quality was evaluated by analyzing read distribution across various genomic regions using bamUtils. Differential gene expression analysis was done with edgeR and DESeq2, followed by Gene Set Enrichment Analysis (GSEA) using the fgsea R package with specified parameters. Curated gene sets from MSigDB were used, and differentially expressed genes were analyzed using DAVID Functional Annotation software with a fold change threshold of <1.2 and FDR <0.05

### Statistics

All data are shown as mean ± SEM. Student t-tests were performed for statistical analysis between the two groups. One-way ANOVA was used to compare three or more groups with a single control group. Statistical analysis was performed using Microsoft Excel and GraphPad Prism 10 (GraphPad Software, San Diego, CA). A P value of 0.05 or less was considered significant.

## Data Availability Statement

All datasets generated during this study are available from the corresponding author (Pratibha Singh) upon reasonable request. Additionally, RNA sequencing data will be deposited in GEO.

## Authors’ Contributions

D.K. performed the experiments, analyzed the data, and participated in interpreting the results. J.R. analyzed and interpreted the RNA sequencing data. M.K. collected, analyzed, and interpreted the confocal imaging data. Y.Q. and N.I. maintained the mouse colonies and assisted with both *in vivo* and *ex vivo* mouse studies. P.S. conceptualized and supervised the overall study, designed and performed the experiments, analyzed and interpreted the data, and wrote the manuscript. All the authors reviewed and approved the manuscript.

## Acknowledgments

We thank the Indiana University (IU) Center for Medical Genomics, the IU Cooperative Center of Excellence in Hematology (CCEH) Flow Cytometry Core, the Experimental Mouse Resources Core (EMRC), the IU Histology Core, and the LARC facility.

