## Supplemental for "Neuropeptide Y deficiency in the bone marrow drives hematopoietic stem and progenitor cell aging"

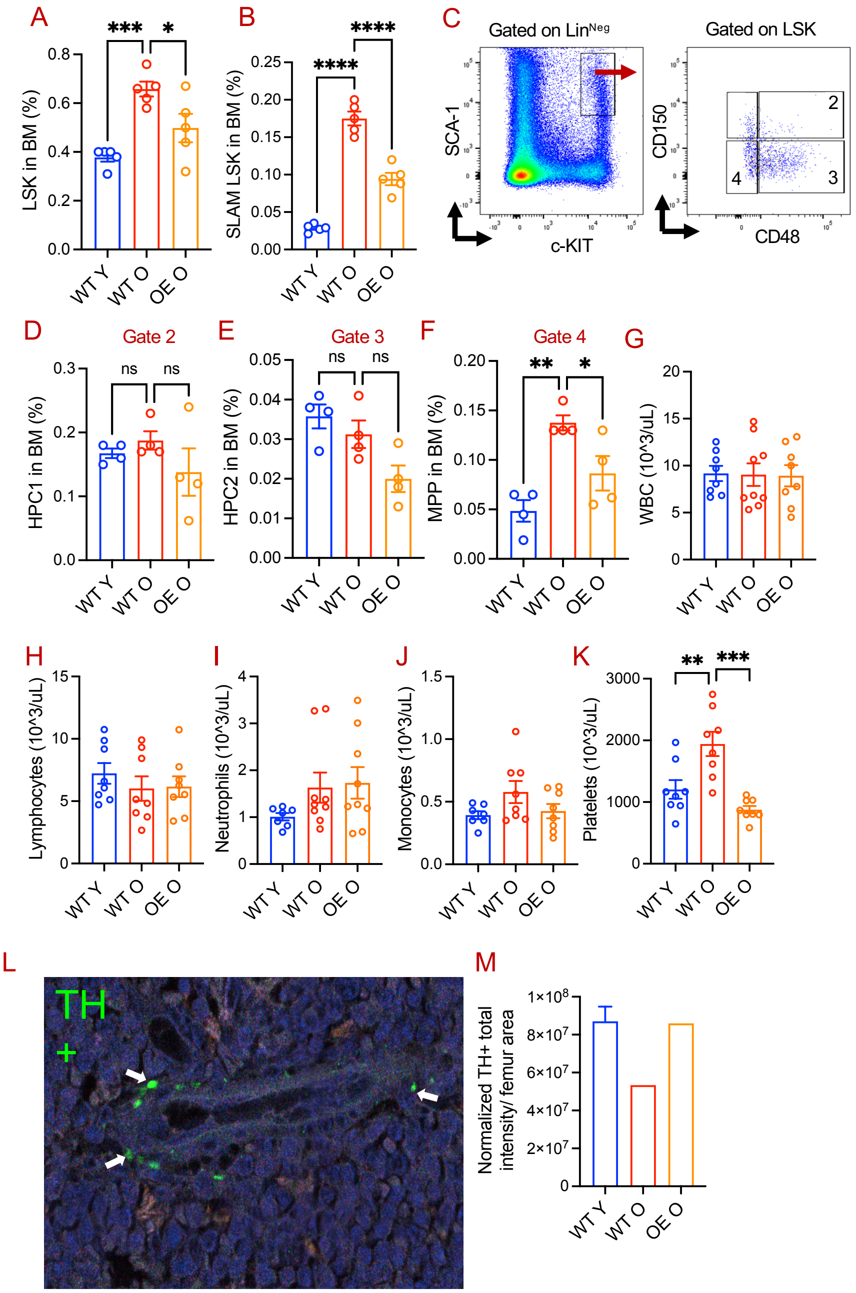
A
LSK in BM (%)
***
*
WT Y
WT O
OE O
B
SLAM LSK in BM (%)
****
****
C
Gated on LinNeg
SCA-1
c-KIT
Gated on LSK
CD150
CD48
2
3
4
D
Gate 2
HPC1 in BM (%)
ns
E
Gate 3
HPC2 in BM (%)
F
Gate 4
MPP in BM (%)
**
G
WBC (10^3/uL)
H
Lymphocytes (10^3/uL)
I
Neutrophils (10^3/uL)
J
Monocytes (10^3/uL)
K
Platelets (10^3/uL)
L
TH
+
M
Normalized TH+ total intensity/ femur area

Supplementary Figure 2

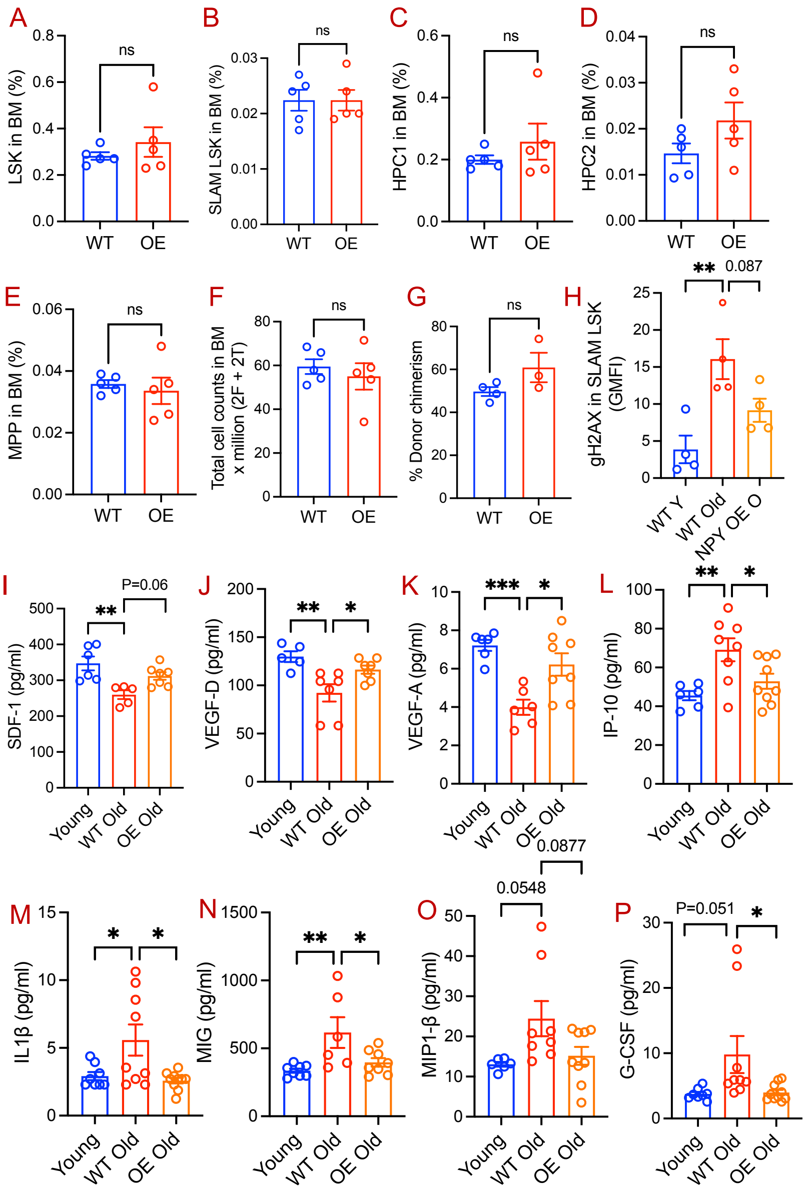

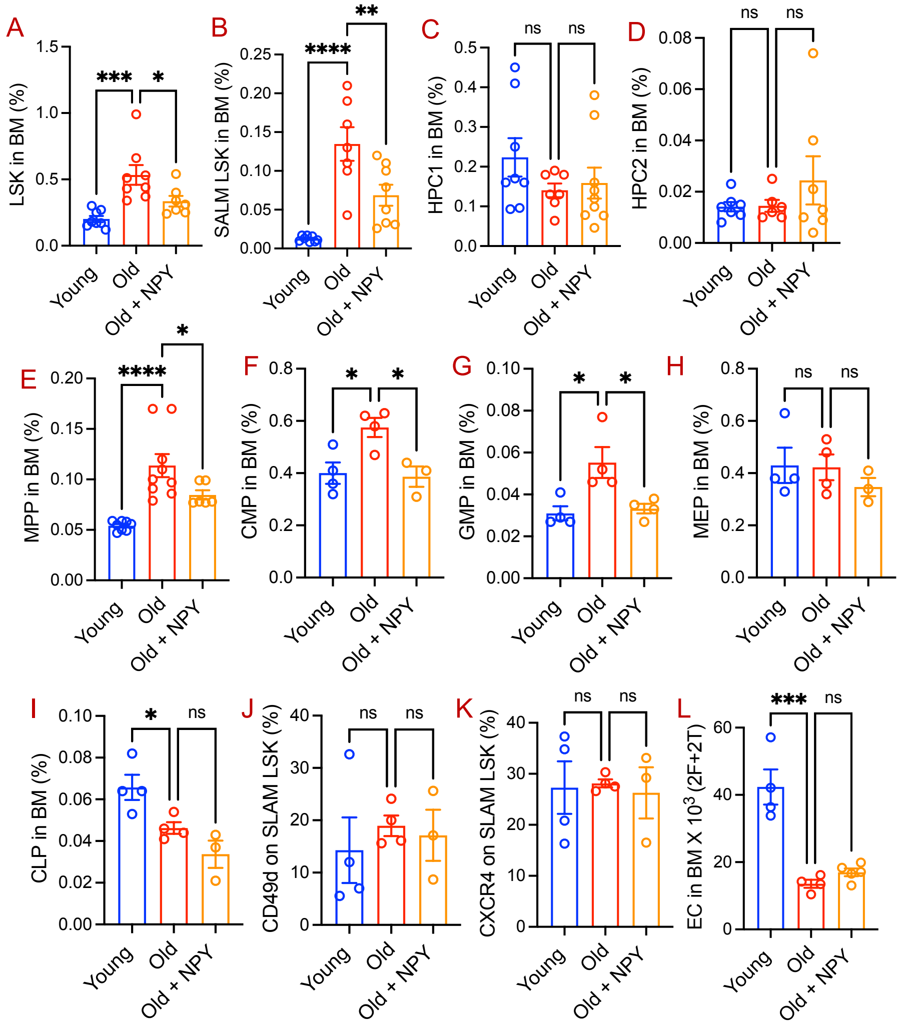
A
LSK in BM (%)
***
*
B
SALM LSK in BM (%)
****
**
C
HPC1 in BM (%)
ns
ns
D
HPC2 in BM (%)
ns
ns
E
MPP in BM (%)
****
*
F
CMP in BM (%)
*
*
G
GMP in BM (%)
*
*
H
MEP in BM (%)
ns
ns
I
CLP in BM (%)
*
ns
J
CD49d on SLAM LSK (%)
ns
ns
K
CXCR4 on SLAM LSK (%)
ns
ns
L
EC in BM X 10^3 (2F+2T)
***
ns
Young
Old
Old + NPY

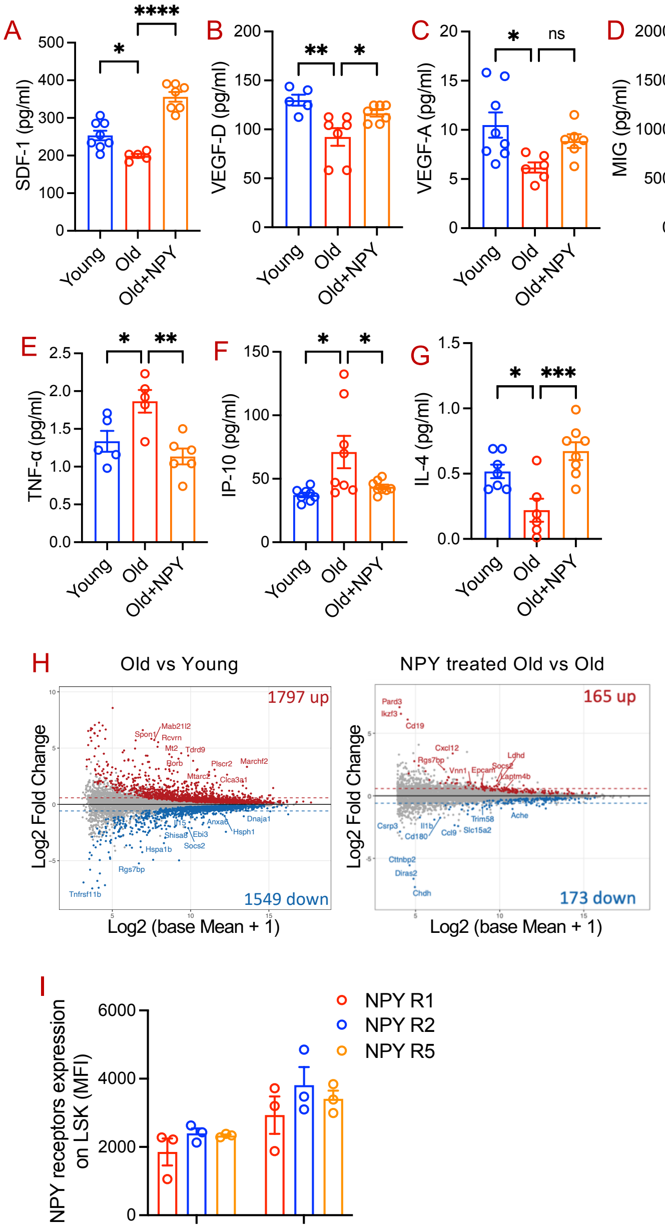

A
SDF-1 (pg/ml)
B
VEGF-D (pg/ml)
C
VEGF-A (pg/ml)
D
MIG (pg/ml)
E
TNF-α (pg/ml)
F
IP-10 (pg/ml)
G
IL-4 (pg/ml)
Young
Old
Old+NPY
H
Old vs Young
NPY treated Old vs Old
1797 up
1549 down
165 up
173 down
Log2 Fold Change
Log2 (base Mean + 1)
I
NPY receptors expression on LSK (MFI)
NPY R1
NPY R2
NPY R5

Supplementary Figure 5

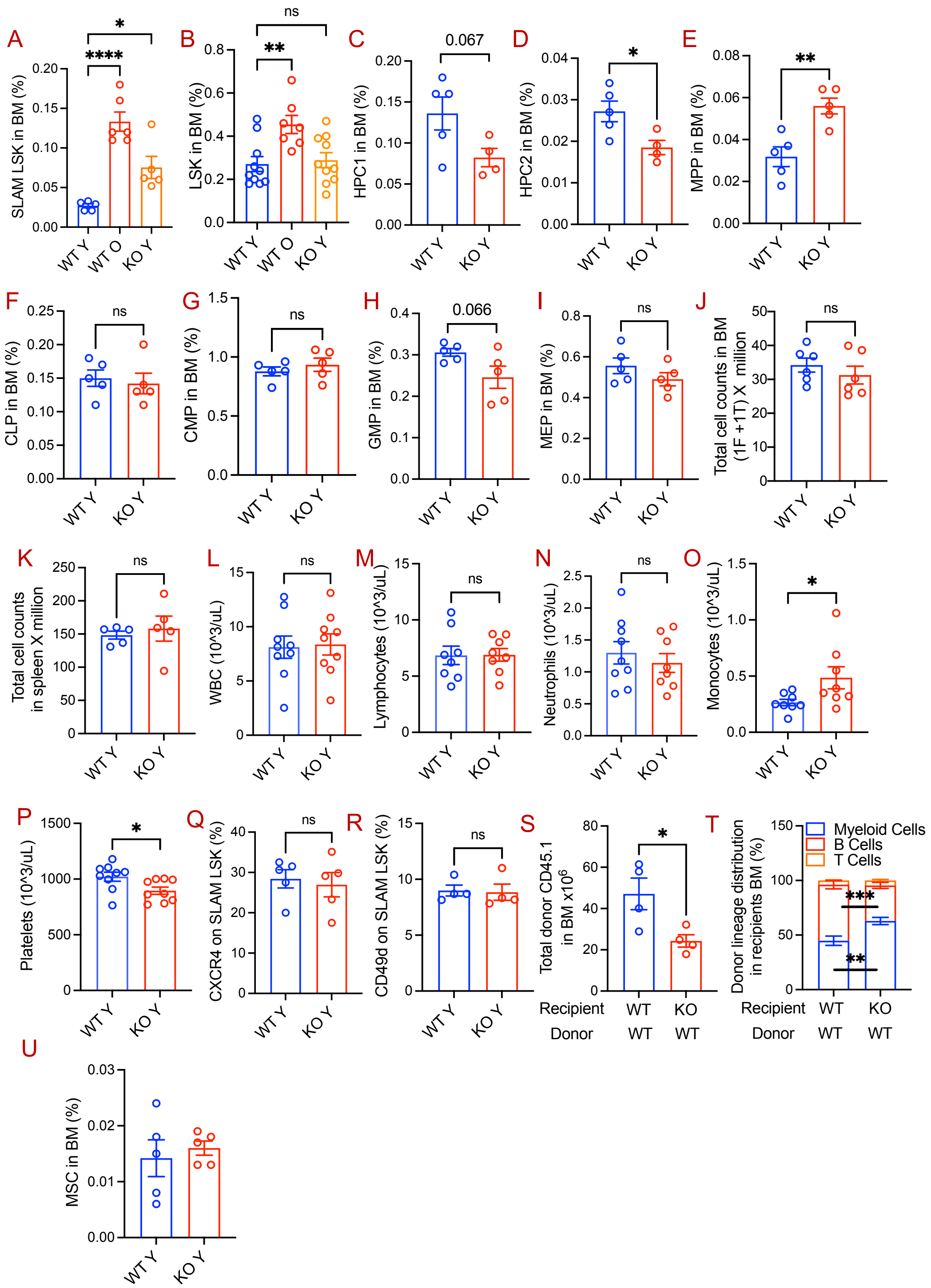

### Supplementary Materials and Methods

**Bone marrow tyrosine hydroxylase staining and imaging:** Mouse femurs were fixed with 4% paraformaldehyde for 48 hours. Following fixation, femurs were decalcified and embedded in paraffin and cut into 5-micron sections to evaluate nerve integrity. Femur sections were stained with anti-mouse tyrosine hydroxylase antibody (Abcam; 1:500 dilution), followed by AF488-conjugated secondary antibody staining. Imaging of femur sections was conducted using a Leica TCS SP8 confocal/2-photon microscope (Leica Microsystems, Inc., Buffalo Grove, IL) at the ICBM Imaging Facility in Indianapolis, IN. Images were captured at a 12-bit depth, with a resolution of 512×512 pixels and a zoom factor 1 (covering an area of 554×554 μm per field). Scanning was performed at a speed of 400 Hz, with a pinhole size of 1 AU, bidirectional X-scanning, and line averaging set to 2. Femur bone cross-section images were analyzed for total tyrosine hydroxylase (TH) signal using Imaris software (version 10.2, Oxford Instruments), available at the Indiana Center for Biological Microscopy (Indianapolis, IN). The nuclei signal was first used to define the bone marrow (BM) area in which the TH signal was subsequently quantified. Whole-tissue nuclei were segmented using the 'Surfaces' module and filtered by axial length. The segmentation results were then manually corrected to remove remaining nuclei from the bone, muscle, disconnected tissue fragments, artifacts, and fatty areas, as these could interfere with the TH readout. The 'Surfaces' module was also used to segment the TH signal in the green channel (488 nm). To isolate the specific TH fluorescence, an arithmetic processing function was applied to subtract the endogenous fluorescence detected in the 552 nm channel from the 488 nm channel. This correction minimized interference from widespread endogenous fluorescence that contaminated the green channel, ensuring accurate detection of TH-specific signal across the BM section. TH-positive surfaces were further filtered based on their distance from BM nuclei and size to ensure that only BM-specific TH signals were included in the analysis, eliminating residual endogenous fluorescence. The accuracy of the analysis method and signal filtering was validated using a negatively stained control tissue. Statistical parameters describing the total area of TH-positive surfaces and BM nuclei per tissue section were exported, and the total TH-positive area was normalized to the BM nuclei area.

**Complete blood cell quantitation:** Peripheral blood was collected from mice by cardiac puncture after CO2 asphyxiation using an EDTA-rinsed syringe. The blood was transferred to EDTA-coated tubes for complete blood cell (CBC) analysis. The CBC with differentials analysis was performed using a Heska Element HT5 Hematology Analyzer.

**Multiplex analysis:** Bone marrow extracellular fluid (BMEF) from the femurs and tibias of mice was collected by crushing them in 1 ml of cold PBS. Bone marrow cell suspensions were centrifuged at 800 g for 5 minutes at 4 °C to collect the supernatants. These supernatants were used to quantify HSC-supporting factors and inflammatory cytokines using Luminex xMAP technology, as we described. Cytokines and HSC-supporting factors were detected using the MILLIPLEX® Mouse Cytokine/Chemokine Magnetic Bead Panel (Millipore, Billerica, MA). Samples were analyzed using the Bio-Plex™ 200 System equipped with High Throughput Fluidics (HTF) Multiplex Array System (Bio-Rad® Laboratories, Hercules, CA). Data analysis was performed using the Bio-Plex 6.0 Manager software from Bio-Rad.

### Supplementary Figure Legends

**Figure S1. Effects of NPY overexpression on aged mice hematopoietic cells and BM sympathetic nerve fibers.** (A-B) LSK and SLAM LSK frequencies in the BM of WT young and old, and NPY OE old mice (C-F). Representative gating strategies of hematopoietic progenitor subpopulations (C), HPC1s (D), HPC2s (E), and MPPs (F) in the BM of WT young and old mice and NPY OE old mice. (G-K) Quantitation of WBC, lymphocytes, neutrophils, monocytes, and platelets by Heska Element HT5 Hematology Analyzer in the PB of WT young, WT old, and NPY OE old mice. (L-M) Evaluation of nerve integrity in the femur sections of WT young and old and NPY OE old mice: (L) A representative image of tyrosine hydroxylase (TH) staining, and (M) Quantitation of TH expression. PFA-fixed mouse femur sections were stained with anti-TH antibody, and TH intensity in these tissue sections was evaluated by confocal microscopy. Data are mean ± S.E.M.; $^{*}p \leq 0.05$, $^{**}p \leq 0.01$, $^{***}p \leq 0.001$).

**Figure S2. Evaluation of HSPCs in young WT and NPY OE mice and age-associated changes in BM niche factors.** (A-E) Flow cytometric quantification of BM HSPC populations in young WT and NPY OE mice: (A) LSK, (B) SLAM LSK, (C) HPC1, (D) HPC2, and (E) MPP. (F) Total BM cellularity in young WT and NPY OE mice. (G) Hematopoietic repopulating capacity of BM cells from young WT and NPY OE mice assessed by competitive transplantation at 3 months post-transplant. (H) γH2AX expression in BM SLAM LSK cells from young and old WT mice and old NPY OE mice. (I-K) HSC-supporting factors and (L-P) inflammatory factors measured in BM extracellular fluid from young and old WT mice and old NPY OE mice by Multiplex assays. Data are mean ± S.E.M.; $^{*}p \leq 0.05$, $^{**}p \leq 0.01$, $^{***}p \leq 0.001$.

**Figure S3. Effects of exogenous NPY administration on old mice HSPCs.** (A-I) Evaluation of HSPCs frequencies by flow cytometry in the BM of young, old and NPY-treated old (100 ng/day; 15 days) mice: (A) LSK, (B) SLAM LSK, (C) HPC1, (D) HPC2, (E) MMP, (F) CMP, (G) GMP, (H) MEP, and (I) CLP. (J-K) Evaluation of cell surface CD49d and CXCR4 expression on the BM SLAM LSK of young, old and NPY treated old mice. (L) ECs is the BM of young, old and NPY treated old mice. Data are mean ± S.E.M.; $^{*}p \leq 0.05$, $^{**}p \leq 0.01$, $^{***}p \leq 0.001$.

**Figure S4. Effects of exogenous NPY administration on the levels of BM factors and transcriptome expression in HSCs of aged mice.** (A-C) Quantitation of HSC supporting factors, and (D-G) inflammatory factors levels by Milliplex Multiplex assays in the BMEF of young, old, and NPY-treated old mice (100 ng/day; 15 days). (H) MA plots showing average expression and fold-change differential gene expression in the HSCs from young, old and NPY treated old mice. Colored points indicate $padj < 0.05$. (I) Evaluation of NPY receptor subtypes (R1, R2, and R5) expression in BM LSK of young and old mice. Data are mean ± S.E.M.; $*p \leq 0.05$, $**p \leq 0.01$).

**Figure S5. Effects of NPY deficiency on hematopoietic cells and MSCs in young mice.** (A-B) Frequencies of LSK and SLAM LSK cells in the BM of young and old WT mice and young NPY KO mice. (C-I) Frequencies of HPC1, HPC2, MPP, CLP, CMP, GMP, and MEP populations in the BM of young WT and NPY KO mice. (J-K) Total cellularity of the BM and spleen in young WT and NPY KO mice. (L-P) Peripheral blood counts of WBCs, lymphocytes, neutrophils, monocytes, and platelets measured using the Heska Element HT5 Hematology Analyzer. (Q-R) CXCR4 and CD49d expression on BM SLAM LSK cells from young WT and NPY KO mice. (S-T) Effects of BM niche NPY deficiency on young WT mice derived donor cells hematopoietic repopulation ability and multilineage differentiation ability. Young WT mice-derived BM cells (one million) were transplanted into lethally irradiated middle-aged (12 months old) WT or NPY KO mice. Three months post-transplantation, donor cells' (S) hematopoietic repopulation, and (T) lineage differentiation were evaluated in the BM of recipient mice. (U)Frequency of BM MSCs ($CD45^-Ter119^-CD31^-CD51^+PDGFR\alpha^+$) in young WT and NPY KO mice assessed by flow cytometry. Data are presented as mean ± S.E.M.; $*p \leq 0.05$, $**p \leq 0.01$.
